# Local online learning in recurrent networks with random feedback

**DOI:** 10.1101/458570

**Authors:** James M. Murray

## Abstract

A longstanding challenge for computational neuroscience has been the development of biologically plausible learning rules for recurrent neural networks (RNNs) enabling the production and processing of time-dependent signals such as those that might drive movement or facilitate working memory. Classic gradient-based algorithms for training RNNs have been available for decades, but they are inconsistent with known biological features of the brain, such as causality and locality. In this work we derive an approximation to gradient-based learning that comports with these biologically motivated constraints. Specifically, the online learning rule for the synaptic weights involves only local information about the pre- and postsynaptic activities, in addition to a random feedback projection of the RNN output error. In addition to providing mathematical arguments for the effectiveness of the new learning rule, we show through simulations that it can be used to train an RNN to successfully perform a variety of tasks. Finally, to overcome the difficulty of training an RNN over a very large number of timesteps, we propose an augmented circuit architecture that allows the RNN to concatenate short-duration patterns into sequences of longer duration.

## Introduction

Many tasks require computations that unfold over time. To accomplish tasks involving motor control, working memory, or other time-dependent phenomena, neural circuits must learn to produce the correct output at the correct time. Such learning is a difficult computational problem, as it generally involves temporal credit assignment, requiring synaptic weight updates at a particular time to minimize errors not only at the time of learning but also at earlier and later times. The problem is also a very general one, as such learning occurs in numerous brain areas and is thought to underlie many complex cognitive and motor tasks encountered in experiments.

To obtain insight into how the brain might perform challenging time-dependent computations, an increasingly common approach is to train high-dimensional dynamical systems known as recurrent neural networks (RNNs) to perform tasks similar to those performed by circuits of the brain, often with the goal of comparing the RNN with neural data to obtain insight about how the brain solves computational problems [1, 2, 3, 4]. While such an approach can lead to useful insights about the neural representations that are formed once a task is learned, it so far cannot address in a satisfying way the process of learning itself, as the standard learning rules for training RNNs suffer from highly nonbiological features such as nonlocality and acausality, as we describe below.

The most straightforward approach to training an RNN to produce a desired output is to define a loss function based on the difference between the RNN output and the target output that we would like it to match, then to update each parameter in the RNN—typically the synaptic weights—by an amount proportional to the gradient of the loss function with respect to that parameter. The most widely used among these algorithms is backpropagation through time (BPTT) [5]. As its name suggests, BPTT is acausal, requiring that errors in the RNN output be accumulated incrementally from the end of a trial to the beginning in order to update synaptic weights. Real-time recurrent learning (RTRL) [6], the other classic gradient-based learning rule, is causal but nonlocal, with the update to a particular synaptic weight in the RNN depending on the full state of the network—a limitation shared by more modern reservoir computing methods [7, 8]. What’s more, both BPTT and RTRL require fine tuning in the sense that the feedback weights from the RNN output back to the network must precisely match the readout weights from the RNN to its output.

The goal of this work is to derive a learning rule for RNNs that is both causal and local, without requiring fine tuning of the feedback weights. Our results depend crucially on two approximations. First, locality is enforced by dropping the nonlocal part of the loss function gradient, making our learning rule only approximately gradient-based. Second, we replace the finely tuned feedback weights required by gradient-based learning with random feedback weights, inspired by the success of a similar approach in nonrecurrent feedforward networks [9, 10]. With these two approximations, we obtain a learning rule that is causal and local, without requiring fine tuning of feedback weights. In the sections that follow, we show that, despite these approximations, RNNs can be effectively trained to perform a variety of tasks. In the Appendices, we provide supplementary mathematical arguments showing why the algorithm remains effective despite its use of an inexact loss function gradient.

## Results

### The RFLO learning rule

To begin, we consider an RNN, as shown in Figure 1, in which a time-dependent input vector **x**(*t*) provides input to a recurrently connected hidden layer of *N* units described by activity vector **h**(*t*), and this activity is read out to form a time-dependent output **y**(*t*). Such a network is defined by the following equations:

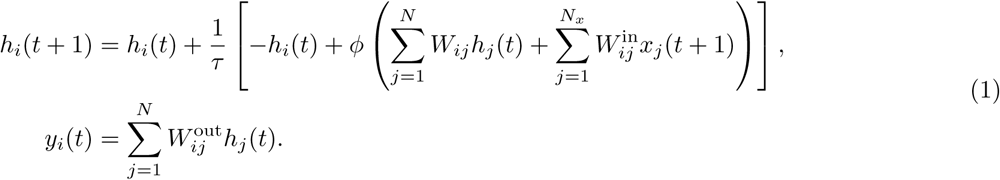

**Figure 1:**
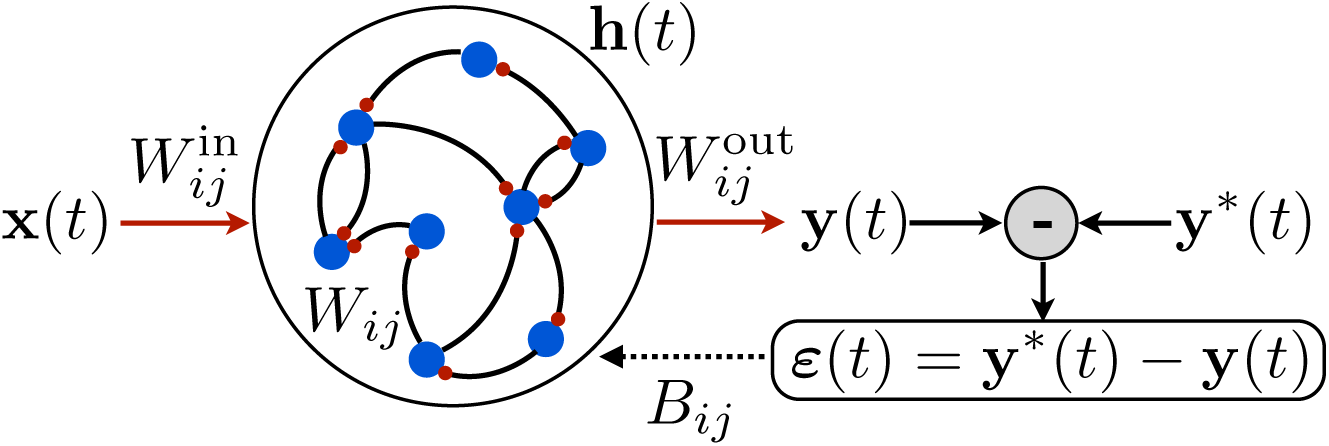
Schematic illustration of a recurrent neural network. The network receives time-dependent input **x**(*t*), and its synaptic weights are trained so that the output **y**(*t*) matches a target function y*(*t*). The projection of the error *ε*(*t*) with feedback weights is used for learning the input weights and recurrent weights.

For concreteness, we take the nonlinear function appearing in (1) to be *ϕ*(·) = tanh(·). The goal is to train this network to produce a target output function **y**\* (*t*) given a specified input function **x**(*t*) and initial activity vector **h**(0). The error is then the difference between the target output and the actual output, and the loss function is the squared error integrated over time:

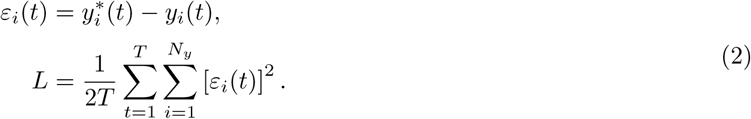

The goal of producing the target output function **y**\*(*t*) is equivalent to minimizing this loss function.

The derivation of the learning rules used in this paper is provided in Appendix 1, where their relation to other gradient-based learning rules (BPTT and RTRL) is also discussed in detail. In brief, the derivation proceeds by taking the gradient of the loss function with respect to the synaptic weights, as in RTRL or BPTT, and then performing gradient descent by updating the synaptic weights by an amount proportional to the gradient. Unlike those methods, however, two approximations are made when taking the gradient. One approximation consists of dropping a nonlocal term from the gradient, so that computing the update to a given synaptic weight requires only pre- and postsynaptic activities, rather than information about the entire state of the RNN including all of its synaptic weights. Second, as described in more detail below, we project the error back into the network for learning using random feedback weights, rather than feedback weights that are tuned to match the readout weights.

Following this procedure, the weight update equations are

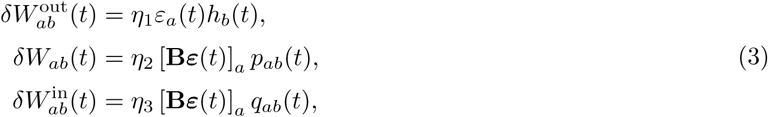

where *η_i_* are learning rates, and **B** is a random matrix of feedback weights. Here we have defined

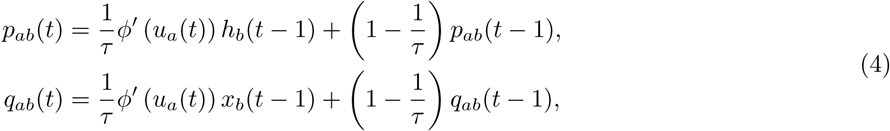

which are the accumulated products of the pre- and (the derivative of the) postsynaptic activity at the recurrent and input synapses, respectively. We have also defined 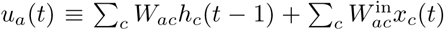 as the total input current to unit *a*. While this form of the update equations does not require explicit integration and hence is more efficient for numerical simulation, it is instructive to take the continuous-time (*τ* ≫ 1) limit of the (14) and the integral of (15), which yields

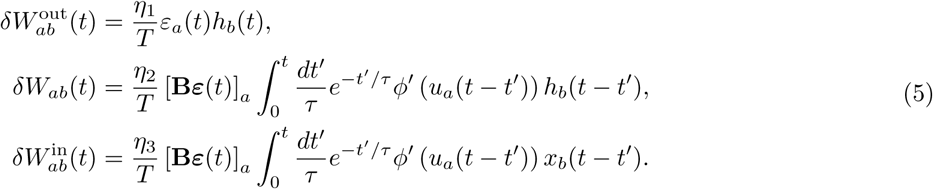

In this way, it becomes clear that the integrals in the second and third equations are *eligibility traces* that accumulate the correlations between pre- and post-synaptic activity over a time window of duration ~ τ. The weight update is then proportional to this eligibility trace, multiplied by a feedback projection of the readout error. The fact that the timescale for the eligibility trace matches the RNN time constant τ reflects the fact that the RNN dynamics are typically correlated only up to this timescale, so that the error is associated only with RNN activity up to time τ in the past. If the error feedback were delayed rather than provided instantaneously, then eligibility traces with longer timescales might be beneficial [11].

Three features of the above learning rules are especially important. First, the updates are local, requiring information about the presynaptic activity and the postsynaptic input current, but no information about synaptic weights and activity levels elsewhere in the network. Second, the updates are online and can either be made at each timestep or accumulated over many timesteps and made at the end of each trial or of several trials. In either case, unlike the BPTT algorithm, it is not necessary to run the dynamics backward in time at the end of each trial to compute the weight updates. Third, the readout error is projected back to each unit in the network with weights **B** that are fixed and random. An exact gradient of the loss function, on the other hand, would lead to (**W**^out^)^T^, where (·)^T^ denotes matrix transpose, appearing in the place of **B**. As described above, the use of random feedback weights is inspired by a similar approach in feedforward networks [9] (see also [12]), and we shall show below that the same feedback alignment mechanism that is responsible for the success of the feedforward version is also at work in our recurrent version.^1^

With the above observations in mind, we refer to the above learning rule as random feedback local online (RFLO) learning. In Appendix 1, we provide a full derivation of the learning rule, and describe in detail its relation to the other gradient-based methods mentioned above, BPTT and RTRL. Because the RFLO learning rule uses an approximation of the loss function gradient rather than the exact gradient for updating the synaptic weights, a natural question to ask is whether it can be expected to decrease the loss function at all. In Appendix 2 we show that, under certain simplifying assumptions including linearization of the RNN, the loss function does indeed decrease on average with each step of RFLO learning. In particular, we show that, as in the feedforward case [9], reduction of the loss function requires alignment between the learned readout weights W^out^ and the fixed feedback weights B. We then proceed to show that this alignment tends to increase during training due to coordinated learning of the recurrent weights **W** and readout weights **W**^out^. Because of the simplifying assumptions made in the mathematical derivations of Appendix 2, however, it remains to be shown that RFLO learning can be used to successfully train a nonlinear RNN in practice. In the following section, therefore, we show using simulated examples that RFLO learning can perform well on a variety of tasks.

### Performance of RFLO learning

In this section we illustrate the performance of the RFLO learning algorithm on a number of simulated tasks. As a benchmark, we compare the performance of RFLO learning with BPTT, the standard algorithm for training RNNs. (As described in Appendix 1, the weight updates in RTRL are, when performed in batches at the end of each trial, completely equivalent to those in BPTT. Hence in this section we compare RFLO learning with BPTT only in what follows.)

### Autonomous production of continuous outputs

Figure 2 illustrates the performance of an RNN trained with RFLO learning to produce a one-dimensional periodic output given no external input. Figure 2a shows the decrease of the loss function (the mean squared error of the RNN output) as the RNN is trained over many trials, where each trial corresponds to one period consisting of *T* timesteps, as well as the performance of the RNN at the end of training. In addition to the RFLO learning rule, BPTT was also used to train the RNN, as well as a variant of RFLO learning (RFLO+Dale) in which all outbound synapses from a given unit were constrained to be of the same sign—a biological constraint known as Dale’s law [13]. In all cases, the sign-constrained network was capable of learning as well as the unconstrained network, as shown in previous modeling work using nonlocal learning rules [14].

**Figure 2:**
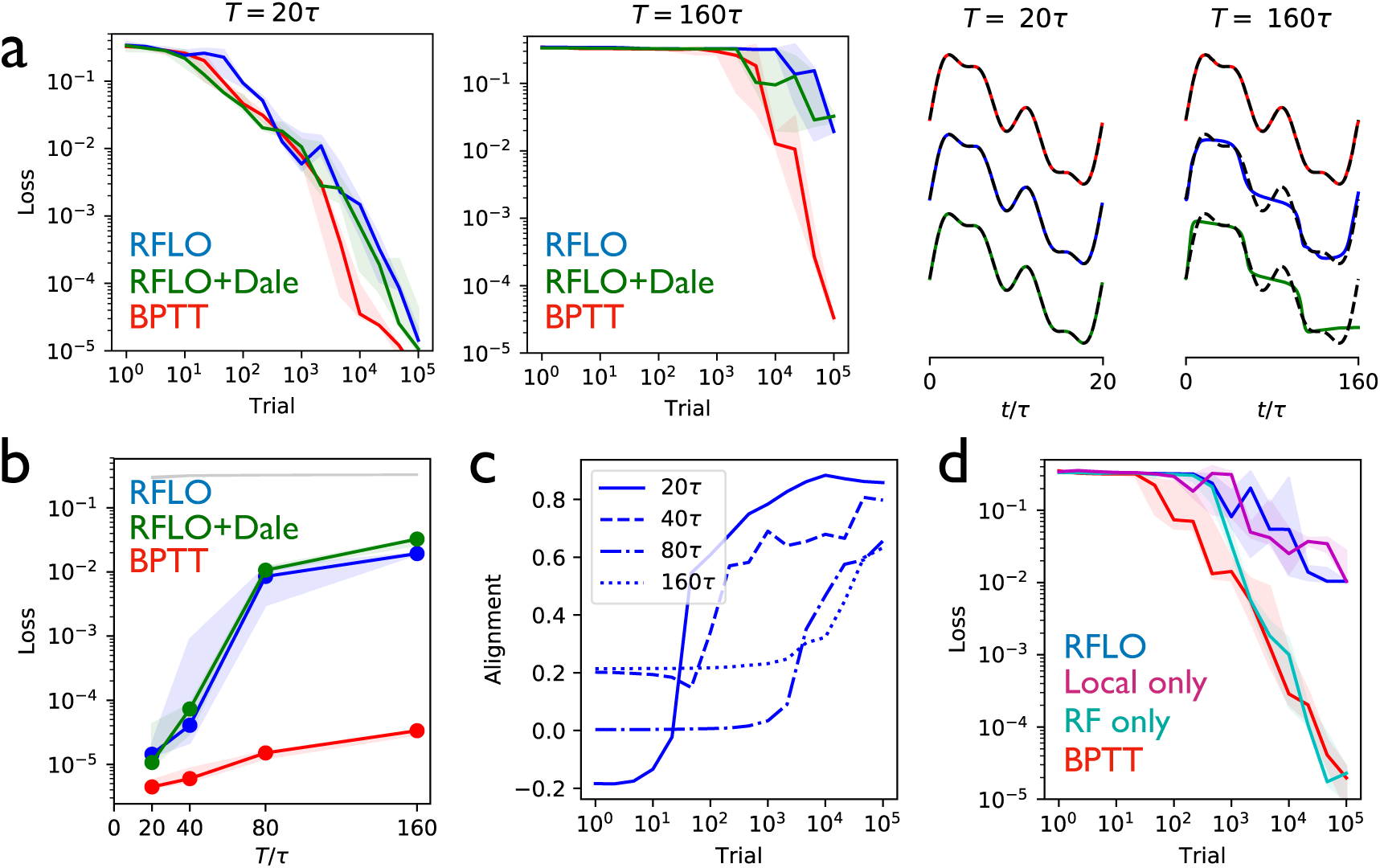
Periodic output task. (a) *Left panels:* The mean squared output error during training for an RNN with *N* = 30 recurrent units and no external input, trained to produce a one-dimensional periodic output with period of duration *T* = 20*τ* (left) or *T* = 160*τ* (right), where *τ* = 10 is the RNN time constant. The learning rules used for training were backpropagation through time (BPTT), RFLO, or RFLO with sign-constrained synapses (RFLO+Dale). *Right panels:* The RNN output at the end of training for each of the three types of learning (dashed lines are target outputs). (b) The loss function at the end of training for target outputs having different periods. The three colors correspond to the three learning rules from (a), while the gray line is the loss computed for an untrained RNN. (c) The normalized alignment between the vector of readout weights **W**^out^ and the vector of feedback weights **B** during training with RFLO learning. (d) The loss function during training with *T* = 80*τ* for BPTT and RFLO, as well as versions of RFLO in which locality is enforced without random feedback (magenta) or random feedback is used without enforcing locality (cyan). In this figure, solid lines show median loss over 5 trained RNNs, learning rates were selected by grid search over *η*_1,2,3_ = *η* € [10^−4^, 10^0^], and the RNN time constant was *τ* = 10.

Figure 2b shows that, in the case where the number of timesteps in the target output was not too great, both versions of RFLO learning perform comparably well to BPTT. BPTT showed an advantage, however, when the number of timesteps became very large. Intuitively, this difference in performance is due to the accumulation of small errors in the estimated gradient of the loss function over many timesteps with RFLO learning. This is less of a problem for BPTT, on the other hand, in which the exact gradient is used.

Figure 2c shows the increase in the alignment between the vector of readout weights **W**^out^ and the vector of feedback weights **B** during training with RFLO learning. As in the case of feedforward networks [9, 12], the readout weights evolve over time to become increasingly similar to the feedback weights, which are fixed during training. In Appendix 2 we provide mathematical arguments for why this alignment occurs, showing that the alignment is not due to the change in **W**^out^ alone, but rather to coordinated changes in the readout and recurrent weights.

In deriving the RFLO learning rule, two independent approximations were made: locality was enforced by dropping the nonlocal term from the loss function gradient, and feedback weights were chosen randomly rather than tuned to match the readout weights. If these approximations are instead made independently, which will have the greater effect on the performance of the RNN? Figure 2d answers this question by comparing RFLO and BPTT with two alternative learning rules: one in which the local approximation is made while symmetric error feedback is maintained, and another in which the nonlocal part of the loss function gradient is retained but the error feedback is random. The results show that the local approximation is essentially fully responsible for the performance difference between RFLO and BPTT, while there is no significant loss in performance due to the random feedback alone.

### Interval matching

Figure 3 illustrates the performance of the RFLO algorithm on a “Ready Set Go” task, in which the RNN is required to produce an output pulse after a time delay matching the delay between two preceding input pulses [15]. This task is more difficult than the production of a periodic output due to the requirement that the RNN must learn to store the information about the interpulse delay, and then produce responses at different times depending on what the delay was. Figures 3b,c illustrate the testing performance of an RNN trained with either RFLO learning or BPTT. If the RNN is trained and tested on interpulse delays satisfying *T*_delay_ ≤ 15*τ*, the performance is similarly good for the two algorithms. If the RNN is trained and tested with longer *T*_delay_, however, then BPTT performs better than RFLO learning. As in the case of the periodic output task from Figure 2, RFLO learning performs well for tasks on short and intermediate timescales, but not as well as BPTT for tasks involving longer timescales. We shall return to this point below in the following subsection.

**Figure 3:**
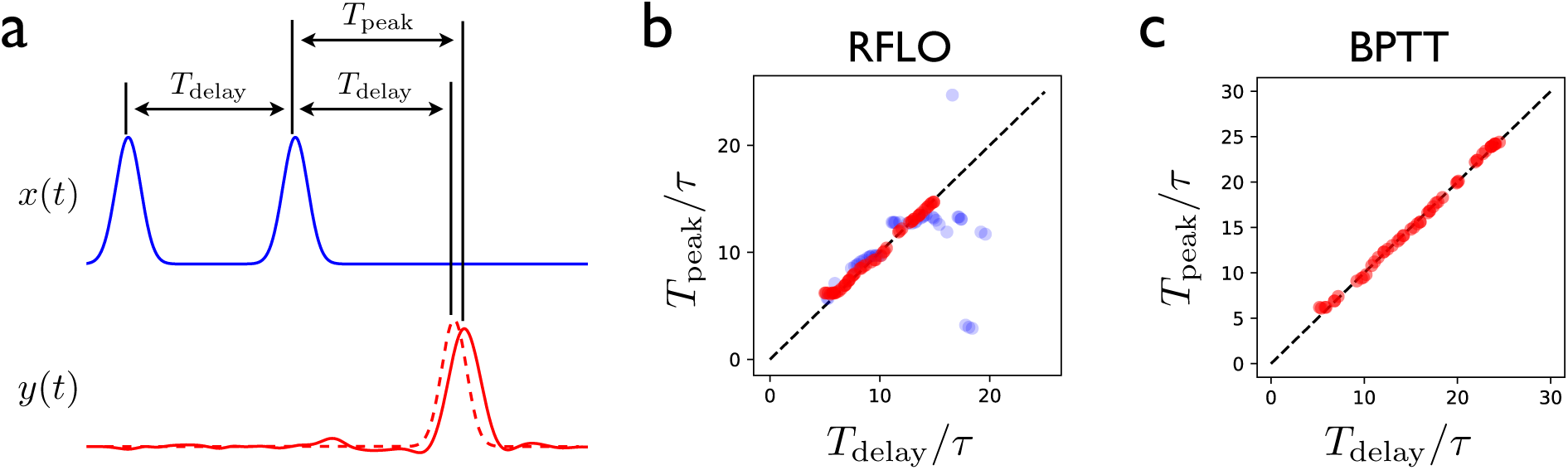
Interval-matching task. (a) In the task, the RNN input consists of two input pulses, with a random delay *T*_delay_ between pulses in each trial. The target output (dashed line) is a pulse trailing the second input pulse by *T*_delay_. (b) The time of the peak in the RNN output is observed after training with RFLO learning and testing in trials with various interpulse delays in the input. Red (blue) shows the case in which the RNN is trained with interpulse delays satisfying *T*_delay_ ≤ 15*τ* (20*τ*). (c) Same as (b), but with the RNN trained using BPTT using interpulse delays *T*_delay_ ≤ 25*τ* for training and testing. The parameters used in all cases were *N* = 100 and *τ* = 10, with 5000 trials of training.

### Learning a sequence of actions

In the above examples, it was shown that, while the performance of RFLO learning is comparable to that of BPTT for tasks over short and intermediate timescales, it is less impressive for tasks involving longer timescales. From the perspective of machine learning, this represents a failure of RFLO learning. From the perspective of neuroscience, however, we can adopt a more constructive attitude. The brain, after all, suffers the same limitations that we have imposed in constructing the RFLO learning rule—namely, causality and locality—and cannot be performing BPTT for learned movements and working memory tasks over long timescales of seconds or more. So how might recurrent circuits in the brain learn to perform tasks over these long timescales? One possibility is that they use a more sophisticated learning rule than the one that we have constructed. While we cannot rule out this possibility, it is worth keeping in mind that, due to the problem of vanishing or exploding gradients, all gradient-based training methods for RNNs fail eventually at long timescales. Another possibility is that a simple, fully connected recurrent circuit in the brain, like an RNN trained with RFLO learning, can only be trained directly with supervised learning over short timescales, and that a more complex circuit architecture is necessary for longer timescales.

It has long been recognized that long-duration behaviors tend to be sequences composed of short, stereotyped actions concatenated together [16]. Further, a great deal of experimental work suggests that learning of this type involves training of synaptic weights from cortex to striatum [17], the input structure of the basal ganglia, which in turn modifies cortical activity via thalamus. In this section we propose a circuit architecture, partially borrowed from Ref. [18] and inspired by the subcortical loop involving basal ganglia and thalamus, that allows an RNN to learn and perform sequences of “behavioral syllables”.

As illustrated in Figure 4a, the first stage of learning in this scheme involves training an RNN to produce a distinct time-dependent output in response to the activation of each of its tonic inputs. In this case, the RNN output is a two-dimensional vector giving the velocity of a cursor moving in a plane. Once the RNN has been trained in this way, the circuit is augmented with a loop structure, shown schematically in Figure 4b. At one end of the loop, the RNN activity is read out with weights *W^s^*. At the other end of the loop, this readout is used to control the input to the RNN. The weights *W^s^* can be learned such that, at the end of one behavioral syllable, the RNN input driving the next syllable in the sequence is activated by the auxiliary loop. This is done most easily by gating the RNN readout so that it can only drive changes at the end of a syllable.

**Figure 4:**
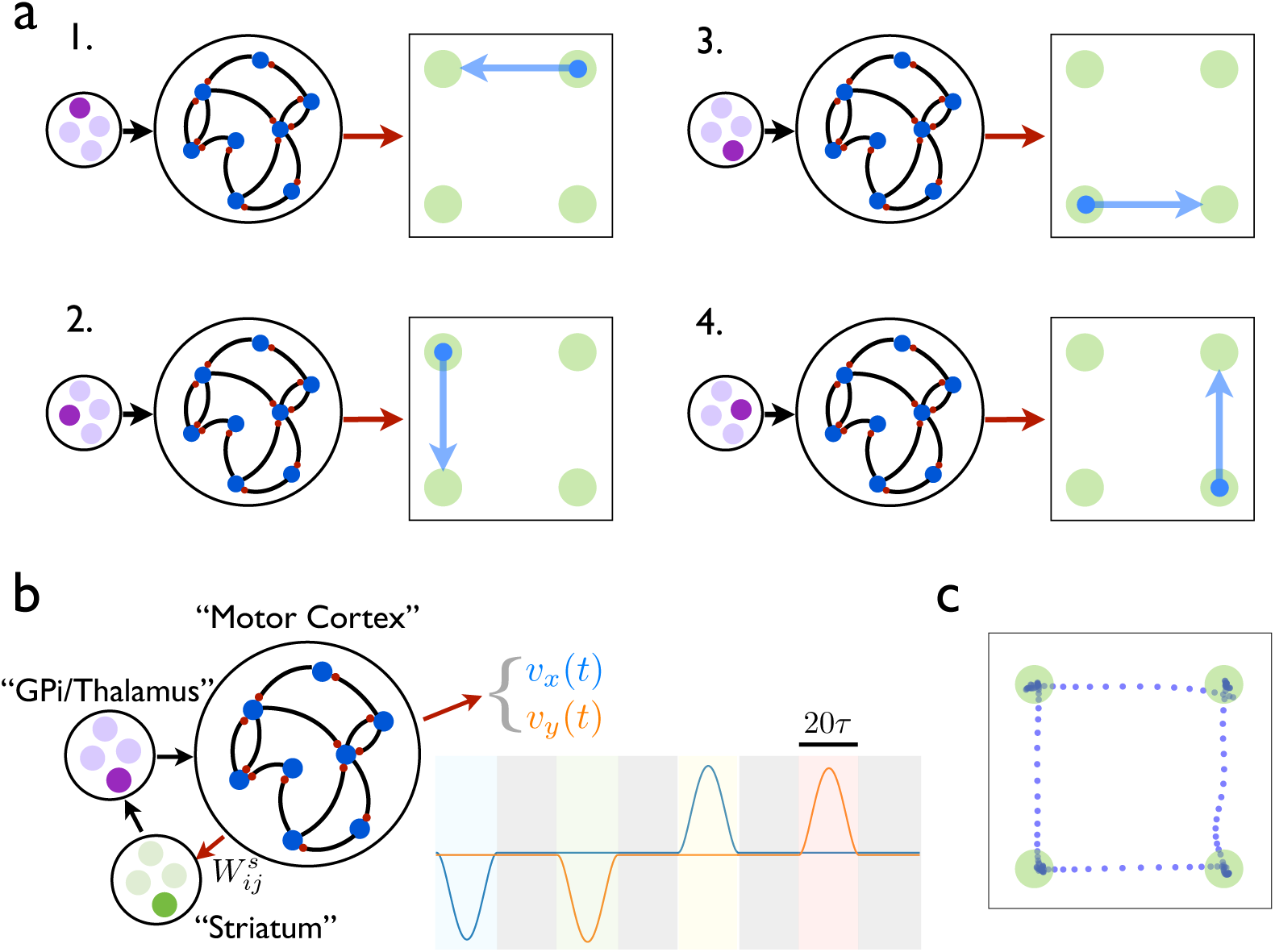
An RNN with multiple inputs controlled by an auxiliary loop learns to produce sequences. (a) An RNN with a two-dimensional readout controlling the velocity of a cursor is trained to move the cursor in a different direction for each of the four possible inputs. (b) The RNN is augmented with a loop structure, which allows a readout from the RNN via learned weights *W^s^* to change the state of the input to the RNN, enabling the RNN state at the end of each cursor movement to trigger the beginning of the next movement. (c) The trajectory of a cursor performing four movements and four holds, where RFLO learning was used to train the individual movements as in (a), and learning of the weights *W^s^* was used to join these movements into a sequence, as illustrated in (b).

In this example, each time the end of a syllable is reached, four readout units receive input 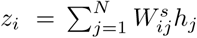, and a winner-take-all rule is applied such that the most active unit activates a corresponding RNN input unit, which drives the RNN to produce the next syllable. Meanwhile, the weights are updated with the reward-modulated Hebbian learning rule 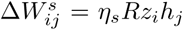, where *R* = 1 if the syllable transition matches the target and *R* = 0 otherwise. By training over many trials, the network learns to match the target sequence of syllables. Figure 4c shows the output from an RNN trained in this way to produce a sequence of reaches and holds in a two-dimensional space. Importantly, while the duration of each behavioral syllable in this example (20*τ*) is relatively short, the full concatenated sequence is long (160*τ*) and would be very difficult to train directly in an RNN lacking such a loop structure.

How might the loop architecture illustrated in Figure 4 be instantiated in the brain? For learned motor control, motor cortex likely plays the role of the recurrent circuit controlling movements. In addition to projections to spinal cord for controlling movement directly, motor cortex also projects to striatum, and experimental evidence has suggested that modification of these corticostriatal synapses plays an important role in the learning of action sequences [19]. Via a loop through the basal ganglia output nucleus GPi and motor thalamus, these signals pass back to motor cortex, as illustrated schematically in Figure 4. According to the model, then, behavioral syllables are stored in motor cortex, and the role of striatum is to direct the switching from one syllable to the next. Experimental evidence for both the existence of behavioral syllables and the role played by striatum in switching between syllables on subsecond timescales has been found recently in mice [20, 21]. How might the weights from motor cortex in this model be gated so that this projection is active at behavioral transitions? It is well known that dopamine, in addition to modulating plasticity at corticostriatal synapses, also modulates the gain of cortical inputs to striatum [22]. Further, it has recently been shown that transient dopamine signals occur at the beginning of each movement in a lever-press sequence in mice [23]. Together, these experimental results support a model in which dopamine bursts enable striatum to direct switching between behavioral syllables, thereby allowing for learned behavioral sequences to occur over long timescales by enabling the RNN to control its own input. Within this framework, RFLO learning provides a biologically plausible means by which the behavioral syllables making up these sequences might be learned.

## Discussion

In this work we have derived an approximation to gradient-based learning rules for RNNs, yielding a learning rule that is local, online, and does not require fine tuning of feedback weights. We have shown that RFLO learning performs comparably well to BPTT when the duration of the task being trained is not too long, but that it performs less well when the task duration becomes very long. In this case, however, we showed that training can still be effective if the RNN architecture is augmented to enable the concatenation of short-duration outputs into longer output sequences. Further exploring how this augmented architecture might map onto cortical and subcortical circuits in the brain is an interesting direction for future work.

How might RFLO learning be implemented concretely in the brain? As we have discussed above, motor cortex is an example of a recurrent circuit that can be trained to produce a particular time-dependent output. Neurons in motor cortex receive information about planned actions (**y**\*(*t*) in the language of the model) from premotor cortical areas, as well as information about the current state of the body (**y**(*t*)) from visual and/or proprioceptive inputs, giving them the information necessary to compute a time-dependent error *ε*(*t*) = **y**\*(*t*) — y(*t*). Hence it is possible that neurons within motor cortex might use a projection of this error signal to learn to produce a target output trajectory. Such a computation might feature a special role for apical dendrites, as in recently developed theories for learning in feedforward cortical networks [31, 32], though further work would be needed to build a detailed theory for its implementation in recurrent cortical circuits.

A possible alternative scenario is that neuromodulators might encode error signals. In particular, midbrain dopamine neurons project to many frontal cortical areas including prefrontal cortex and motor cortex, and their input is known to be necessary for learning certain time-dependent behaviors [33, 34]. Further, recent experiments have shown that the signals encoded by dopamine neurons are significantly richer than the reward prediction error that has traditionally been associated with dopamine, and include phasic modulation during movements [35, 23, 36]. This interpretation of dopamine as a continuous online error signal used for supervised learning would be distinct from and complementary to its well known role as an encoder of reward prediction error for reinforcement learning.

In addition to RFLO learning, a number of other local and causal learning rules for training RNNs have been proposed. The oldest of these algorithms [24, 25] operate within the framework of reinforcement learning rather than supervised learning, meaning that only a scalar—and possibly temporally delayed— reward signal is available for training the RNN, rather than the full target function *y*\*(*t*). Typical of such algorithms, which are often known as “node perturbation” algorithms, is the REINFORCE learning rule [25], which in our notation gives the following weight update at the end of each trial:

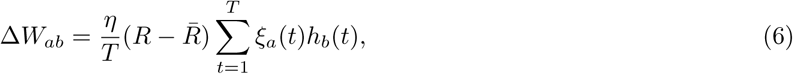

where *R* is the scalar reward signal (which might be defined as the negative of the loss function that we have used in RFLO learning), *R̅* is the average reward over recent trials, and *ξ_a_*(*t*) is noise current injected into unit *a* during training. This learning rule means, for example, that (assuming the presynaptic unit *b* is active) if the postsynaptic unit *a* is more active than usual in a given trial (i.e. *ξ_a_*(*t*) is positive) and the reward is greater than expected, then the synaptic weight *W_ab_* should be increased so that this postsynaptic unit should be more active in future trials. A slightly more elaborate version of this learning rule replaces the summand in (6) with a low-pass filtered version of this same quantity, leading to eligibility traces of similar form to those appearing in (5). Fiete and Seung [26] have developed a version of this learning rule for spiking neurons.

A potential shortcoming of the REINFORCE learning rule is that it depends on the postsynaptic noise current rather than on the total postsynaptic input current (i.e. the noise current plus the input current from presynaptic units). Because it is arguably implausible that a neuron could keep track of these sources of input current separately, a recently proposed version [27] replaces 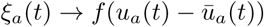, where *f* (·) is a supralinear function, *u_a_* (*t*) is the total input current (including noise) to unit *a*, and *u̅_a_*(*t*) is the low-pass-filtered input current. This substitution is logical since the quantity *u_a_*(*t*) — *u̅_a_*(*t*) tracks the fast fluctuations of each unit, which are mainly due to the rapidly fluctuating input noise rather than to the more slowly varying recurrent and feedforward inputs.

A severe limitation of reinforcement learning as formulated in (6) is the sparsity of reward information, which comes in the form of a single scalar value at the end of each trial. Clearly this provides the RNN with much less information to learn from than a vector of errors *ε*(*t*) = **y**\*(*t*) — **y**(*t*) at every timestep, which is assumed to be available in supervised learning. As one would expect from this observation, reinforcement learning is typically much slower than supervised learning in RNNs, as in feedforward neural networks. A hybrid approach is to assume that reward information is scalar, as in reinforcement learning, but available at every timestep, as in supervised learning. This might correspond to setting *R*(*t*) Ⅱ —|*ε*(*t*)|^2^ and including this reward in a learning rule such as the REINFORCE rule in (6). To our knowledge this has not been done for training recurrent weights in an RNN, though a similar idea has recently been used for training the readout weights of an RNN [28]. Ultimately, whether recurrent neural circuits in the brain use reinforcement learning or supervised learning is likely to depend on the task being learned and what feedback information about performance is available. For example, in a reach-to-target task such as the one modeled in Figure 4, it is plausible that a human or nonhuman primate might have a mental template of an ideal reach, and might make corrections to make the hand match the target trajectory at each timepoint in the trial. On the other hand, if only delayed binary feedback is provided in an interval-matching task such as the one modeled in Figure 3, neural circuits in the brain might be more likely to use reinforcement learning.

More recently, local, online algorithms for supervised learning in RNNs with spiking neurons have been proposed. Gilra and Gerstner [29] and Alemi et al [30] have trained spiking RNNs to produce particular dynamical trajectories of RNN readouts. These works constitute a large step toward greater biological plausibility, particularly in their use of local learning rules and spiking neurons. Here we describe the most important differences between those works and RFLO learning. In both Ref. [29] and [30], the RNN is driven by an input **x**(*t*) as well as the error signal *ε*(*t*) = **y**\*(*t*) — **y**(*t*), where the target output is related to the input x(*t*) according to

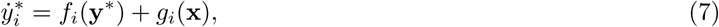

where *g_i_*(**x**) = *x_i_*(*t*) in Ref. [30], but is arbitrary in Ref. [29]. In either case, however, it is not possible to learn arbitrary, time-dependent mappings between inputs and outputs in these networks, since the RNN output must take the form of a dynamical system driven by the RNN input. This is especially limiting if one desires that the RNN dynamics should be autonomous, so that **x**(*t*) = 0 in (7). It is not obvious, for example, what dynamical equations having the form of (7) would provide a solution to the interval-matching task studied in Figure 3. Of course, it is always possible to obtain an arbitrarily complex readout by making x(*t*) sufficiently large such that y(*t*) simply follows x(*t*) from (7). However, since x(*t*) is provided as input, the RNN essentially becomes an autoencoder in this limit.

Two other features of Refs. [29, 30] differ from RFLO learning. First, the readout weights and the error feedback weights are related to one another in a highly specific way, being either symmetric with one another [30], or else configured such that the loop from the RNN to the readout and back to the RNN via the error feedback pathway forms an autoencoder [29]. In either case these weights are preset to these values before training of the RNN begins, unlike the randomly set feedback weights used in RFLO learning. Second, both approaches require that the error signal *ε*(*t*) be fed back to the network with (at least initially) sufficiently large gain such that the RNN dynamics are essentially slaved to produce the target readout **y**\*(*t*), so that one has y(*t*) ≈ **y**\*(*t*) immediately from the beginning of training. (This follows as a consequence of the relation between the readout and feedback weights described above.) With RFLO learning, in contrast, forcing the output to always follow the target in this way is not necessary, and learning can work even if the RNN dynamics early in learning do not resemble the dynamics of the ultimate solution.

In summary, the random feedback learning rule that we propose offers a potential advantage over previous biologically plausible learning rules by making use of the full time-dependent, possibly multidimensional error signal, and also by training all weights in the network, including input, output, and recurrent weights. In addition, it does not require any special relation between the RNN inputs and outputs, nor any special relationship between the readout and feedback weights, nor a mechanism that restricts the RNN dynamics to always match the target from the start of training. Especially when extended to allow for sequence learning such as depicted in Figure 4, RFLO learning provides a plausible mechanism by which supervised learning might be implemented in recurrent circuits in the brain.

## Acknowledgments

The author is grateful to L.F. Abbott, G.S. Escola, and A. Litwin-Kumar for helpful discussions and feedback on the manuscript. Support for this work was provided by the National Science Foundation NeuroNex program (DBI-1707398), the National Institutes of Health (DP5 OD019897), and the Gatsby Charitable Foundation.

## Footnotes

1 While an RNN is often described as being “unrolled in time“, so that it becomes a feedforward network in which each layer corresponds to one timestep, it is important to note that the unrolled version of the problem that we consider here is not identical to the feedforward case considered in Refs. [9, 12]. In the RNN, a readout error is defined at every “layer” *t*, whereas in the feedforward case, the error is defined only at the last layer (*t* = *T*) and is fed back to update weights in all preceding layers.

2 Thanks to A. Litwin-Kumar for discussion about this correspondence.

## Appendix 1 Gradient-based RNN learning and RFLO learning

In the first subsection of this appendix, we begin by reviewing the derivation of RTRL [25], the classic gradient-based learning rule. We show that the update equation for the recurrent weights under the RTRL rule has two undesirable features from a biological point of view. First, the learning rule is nonlocal, with the update to weight *W_ij_* depending on all of the other weights in the RNN, rather than just on information that is locally available to that particular synapse. Second, the RTRL learning rule requires that the error in the RNN readout be fed back into the RNN with weights that are precisely symmetric with the readout weights. In the second subsection, we implement approximations to the RTRL gradient in order to overcome these undesirable features, leading to the RFLO learning rules.

In the third subsection of this appendix, we review the derivation of BPTT, the most widely used algorithm for training RNNs. Because it is the standard gradient-based learning rule for RNN training, BPTT is the learning rule against which we compare RFLO learning in the main text. Finally, in the final subsection of this appendix we illustrate the equivalence of RTRL and BPTT. Although this is not strictly necessary for any of the results given in the main text, we expect that readers with an interest in gradient-based learning rules for training RNNs will be interested in this correspondence, which to our knowledge has not been very clearly explicated in the literature.

### Real-time recurrent learning

In this section we review the derivation of the real-time recurrent learning (RTRL) algorithm [6] for an RNN such as the one shown in Figure 1. This rule is obtained by taking a gradient of the mean-squared output error of the RNN with respect to the synaptic weights, and, as we will show later in this appendix, is equivalent (when implemented in batches rather than online) to the more widely used backpropagation through time (BPTT) algorithm.

The standard RTRL algorithm is obtained by calculating the gradient of the loss function (2) with respect to the RNN weights, and then using gradient descent to find the weights that minimize the loss function [37]. Specifically, for each run of the network, one can calculate *∂L*/*∂W_ab_* and then update the weights by an amount proportional to this gradient: Δ*W_ab_* = —*η∂L*/*∂W_ab_*, where *η* determines the learning rate. This can be done similarly for the input and output weights, 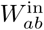 and 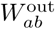, respectively. This results in the following update equations:

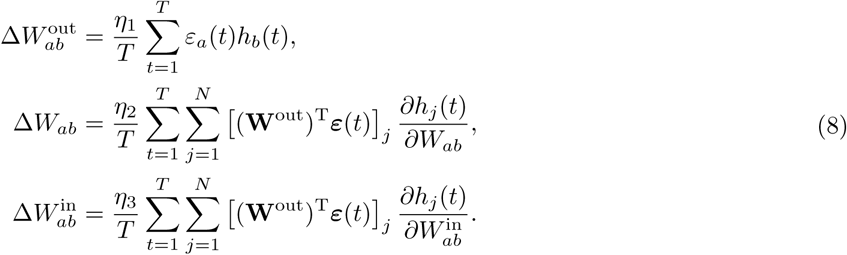

In these equations, (·)^T^ denotes matrix transpose, and the gradients of the hidden layer activities with respect to the recurrent and input weights are given by

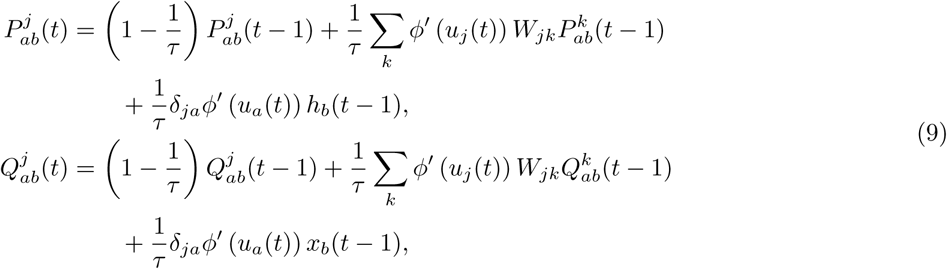

where we have defined

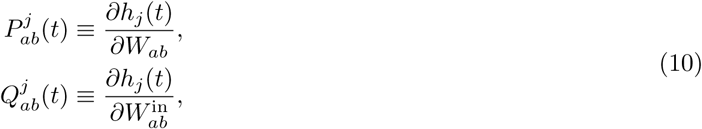

and **u**(*t*) is the total input to each recurrent unit at time *t*:

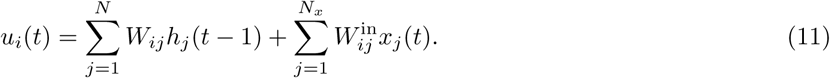

The recursions in (9) terminate with

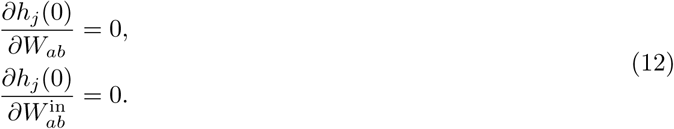

As many others have recognized previously, the synaptic weight updates given in the second and third lines of (8) are not biologically realistic for a number of reasons. First, the error is projected back into the network with the particular weight matrix (*W*^out^)^T^, so that the feedback and readout weights must be related to one another in a highly specific way. Second, the terms involving **W** in (9) mean that information about the entire network is required to update any given synaptic weight, making the rules nonlocal. In contrast, a biologically plausible learning rule for updating a weight W_ab_ or 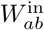 ought to depend only on the activity levels of the pre- and post-synaptic units *a* and *b*, in addition to the error signal that is fed back into the network. Both of these shortcomings will be addressed in the following subsection.

### Random feedback local online learning

In order to obtain a biologically plausible learning rule, we can attempt to relax some of the requirements in the RTRL learning rule and see whether the RNN is still able to learn effectively. Inspired by a recently used approach in feedforward networks [9], we do this by replacing the (*W*^out^)^T^ appearing in the second and third lines of (8) with a fixed random matrix **B**, so that the feedback projection of the output error no longer needs to be tuned to match the other weights in the network in a precise way. Second, we simply drop the terms involving **W** in (9), so that nonlocal information about all recurrent weights in the network is no longer required to update a particular synaptic weight. In this case we can rewrite the approximate weight-update equations as

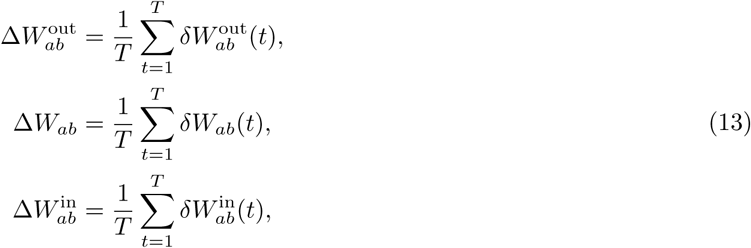

where

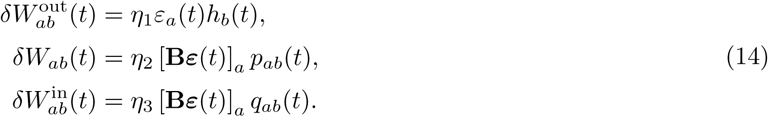

Here we have defined rank-2 versions of the eligibility trace tensors from (10):

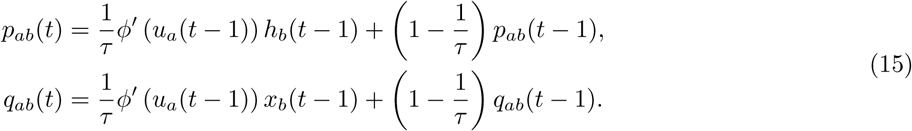

As desired, the equations (13) are local, depending only on the pre- and post-synaptic activity, together with a random feedback projection of the error signal. In addition, because all of the quantities appearing in (13) are computed in real time as the RNN is run, the weight updates can be performed *online*, in contrast to BPTT, for which the dynamics over all timesteps must be run first forward and then backward before making any weight updates. Hence, we refer to the learning rule given by (13)-(10) as random feedback local online (RFLO) learning.

### Backpropagation through time

Because it is the standard algorithm used for training RNNs, in this section we review the derivation of the learning rules for backpropagation through time (BPTT) [5] in order to compare it with the learning rules presented above. The derivation here follows Ref. [38].

Consider the following Lagrangian function:

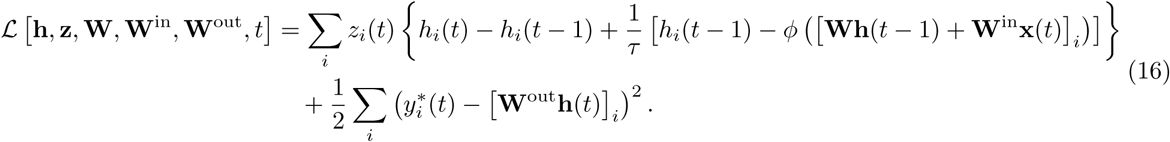

The second line is the cost function that is to be minimized, while the first line uses the Lagrange multiplier z(*t*) to enforce the constraint that the dynamics of the RNN should follow (1). From (16) we can also define the following action:

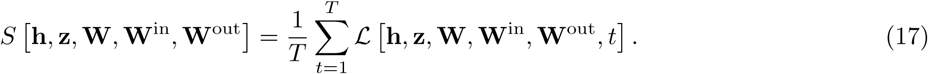

We now proceed by minimizing (17) with respect to each of its arguments. First, taking dS/dz_i_(*t*) just gives the dynamical equation (1). Next, we set *∂S*/*∂h_i_*(*t*) = 0, which yields

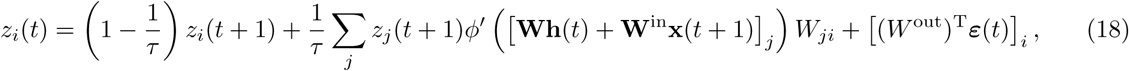

which applies at timesteps *t* = 1,…,*T* — 1. To obtain the value at the final timestep, we take *∂S*/*∂z_i_*(*T*), which leads to

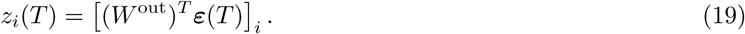

Finally, taking the derivative with respect to the weights leads to the following:

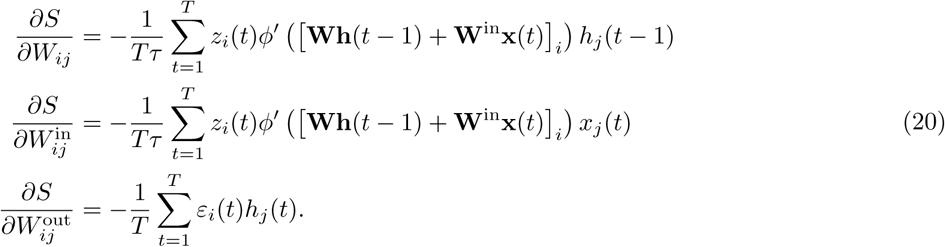

Rather than setting these derivatives equal to zero, which may lead to an undesired solution that corresponds to a maximum or saddle point of the action and would in any case be intractable, we use the gradients in (20) to perform gradient descent, reducing the error in an iterative fashion:

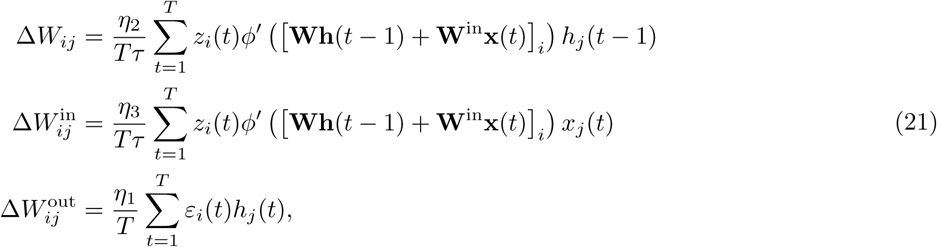

where *η_i_* are learning rates.

The BPTT algorithm then proceeds in three steps. First the dynamical equations (1) for **h**(*t*) are integrated forward in time, beginning with the initial condition **h**(0). Second, the auxiliary variable **z**(*t*) is integrated *backwards* in time using (18), using with the **h**(*t*) saved from the forward pass and the boundary condition **z**(*t*) from (19). Third, the weights are updated according to (21), using **h**(*t*) and **z**(*t*) saved from the preceding two steps.

Note that no approximations have been made in computing the gradients using either the RTRL or BPTT procedures. In fact, as we will show in the following section, the two algorithms are completely equivalent, at least in the case where RFLO weight updates are performed only at the end of each trial rather than at every timestep.

### A unified view of gradient-based learning in recurrent networks

As pointed out previously [39, 40], the difference between RTRL and BPTT can ultimately be traced to distinct methods of bookkeeping in applying the chain rule to the gradient of the loss function.^2^ In order to make this explicit, we begin by noting that, when taking implicit dependences into account, the loss function defined in (2) has the form

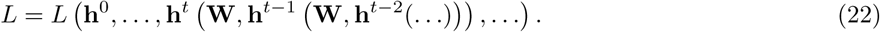

In this section, we write **h**^*t*^ ≡ **h**(*t*) for notational convenience, and consider only updates to the recurrent weights **W**, ignoring the input x(*t*) to the RNN. In any gradient-based learning scheme, the weight update Δ*W_ab_* should be proportional to the gradient of the loss function, which has the form

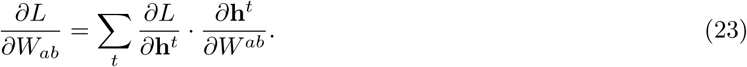

The difference between RTRL and BPTT arises from the two possible ways of keeping track of the implicit dependencies from (22), which give rise to the following equivalent formulations of (23):

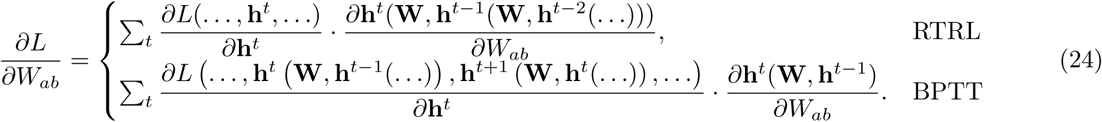

In RTRL, the first derivative is simple to compute because loss function is treated as an explicit function of the variables **h**^*t*^. The dependence of **h**^*t*^ on **W** and **h**^*t′*^ (where *t′* < *t*) is then taken into account in the second derivative, which must be computed recursively due to the nested dependence on **W**. In BPTT, on the other hand, the implicit dependencies are dealt with in the first derivative, which in this case must be computed recursively because all terms at times *t′* > *t* depend implicitly on **h**^*t*^. The second derivative then becomes simple since these dependencies are no longer present.

Let us define the following:

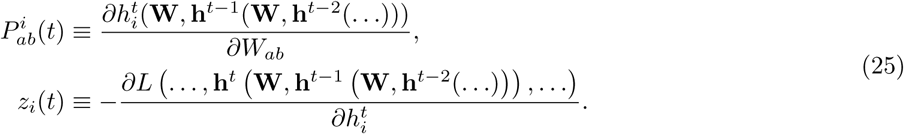

Then, using the definition of *L* from (2) and the dynamical equation (1) for **h**^*t*^ to take the other derivatives appearing in (24), we have

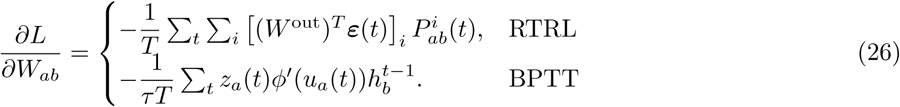

The recursion relations follow from application of the chain rule in the definitions from (25):

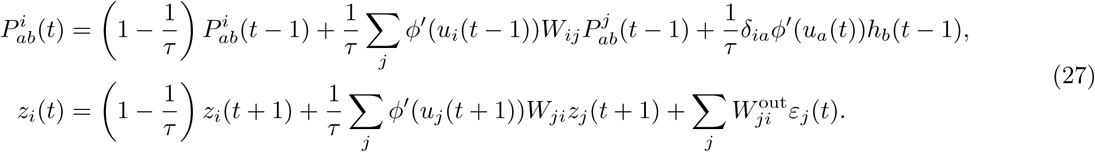

These recursion relations are identical to those appearing in (9) and (18). Notably, the first is computed forward in time, while the second is computed backward in time. Because no approximations have been made in computing the gradient in either case for (26), the two methods are equivalent, at least if RTRL weight updates are made only at the end of each trial, rather than online. For this reason, only one of the algorithms (BPTT) was compared against RFLO learningx in the main text.

As discussed in previous sections, RTRL has the advantages of obeying causality and of allowing for weights to be continuously updated. But, as discussed above, RTRL has the disadvantage of being nonlocal, and also features a greater computational cost due to the necessity of updating a rank-3 tensor 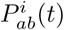 rather than a vector *Z_i_*(*t*) at each timestep. By dropping the second term in the first line of (27), RFLO learning eliminates both of these undesirable features, so that the resulting algorithm is causal, online, local, and has a computational complexity (~ *N*^2^ per timestep, vs. ~ *N*^4^ for RTRL) on par with BPTT.

## Appendix 2 Analysis of the RFLO learning rule

Given that the learning rules in (5) do not move the weights directly along the steepest path that would minimize the loss function (as would the learning rules in (8)), it is worthwhile to ask whether it can be shown that these learning rules in general decrease the loss function at all. To answer this question, we consider the change in weights after one trial lasting T timesteps, working in the continuous-time limit for convenience, and performing weight updates only at the end of the trial:

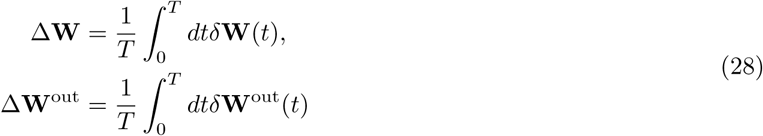

where *δ***W** and *δ***W**^out^ are given by (5). For simplicity in this section we ignore the updates to the input weights, since the results in this case are very similar to those for recurrent weight updates.

In the first subsection of this appendix, we show that, under some approximations, the loss function tends to decrease on average under RFLO learning if there is positive alignment between the readout weights W^out^ and the feedback weights B. In the second subsection, we show that this alignment tends to increase during RFLO learning.

### Decrease of the loss function

We first consider the change in the loss function defined in (2) after updating the weights:

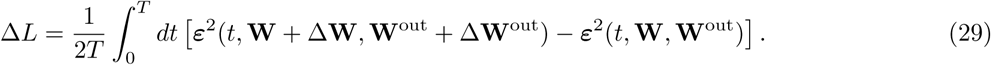

Assuming the weight updates to be small, we ignore terms beyond leading order in ΔW and ΔW^out^. Then, using the update rules in (28) and performing some algebra, (29) becomes

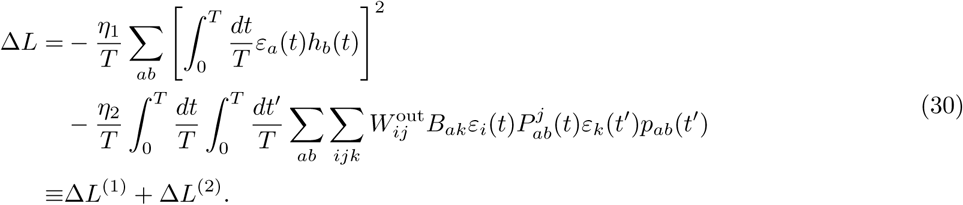

Clearly the first term in (30) always tends to decrease the loss function, as we would expect given that the precise gradient of *L* with respect to **W**^out^ was used to determine this part of the learning rule. We now wish to show that, at least on average and with some simplifying assumptions, the second term in (30) tends to be negative as well. Before beginning, we note in passing that this term is manifestly nonpositive like the first term if we perform RTRL, in which case 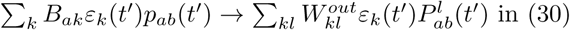, making the gradient exact.

In order to analyze Δ*L*^(2)^, we will assume that the RNN is linear, with *ϕ*(*x*) = *x*. Further, we will average over the RNN activity **h**(*t*), assuming that the activities are correlated from one timestep to the next, but not from one unit to the next:

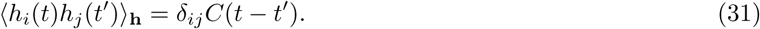

The correlation function should be peaked at a positive value at *t* — *t′* = 0 and decay to 0 at much earlier and later times. Finally, because of the antisymmetry under *x* → —*x*, odd powers of **h** will average to zero: 〈*h_i_*〉_h_ = 〈*h_i_h_j_h_k_*〉_h_ = °.

With these assumptions, we can express the activity-averaged second line of (30) as 〈Δ*L*^(2)^〉_h_ = *F*_1_ + *F*_2_, with

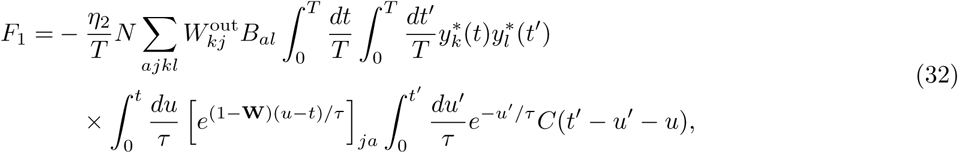

and

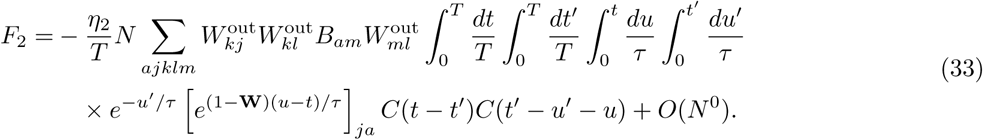

In order to make further progress, we can perform an ensemble average over **W**, assuming that *W_ij_ N*(0, *g*^2^/*N*) is a random variable, which leads to

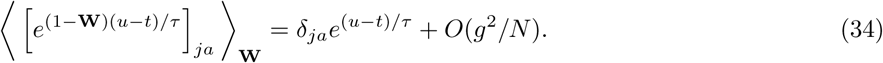

This leads to

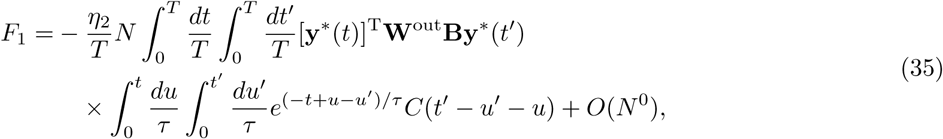

and

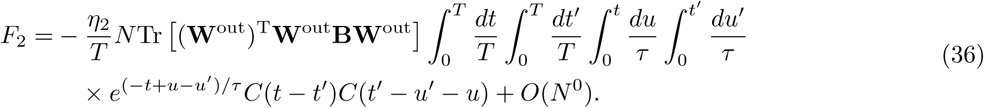

Putting (35) and (36) together, changing one integration variable, and dropping the terms smaller than *O*(*N*) then gives

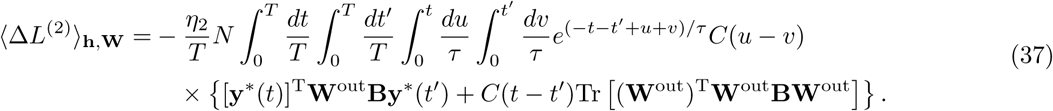

Because we have assumed that *C*(*t*) ≥ 0, the sign of this quantity depends only on the sign of the two terms in the second line of (37).

Already we can see that (37) will tend to be negative when W^out^ is aligned with *B*. To see this, suppose that **B** = *α***W**^out^, with *α* > 0. Due to the exponential factor, the integrand will be vanishingly small except when *t* ≈*t′*, so that the first term in the second line in this case can be written as ≈ *α*|(**W**^out^)^T^**y**\*(*t*)|^2^ ≥ 0. The second term, meanwhile, becomes *αC*(*t* — *t′*)Tr [((**W**^out^)^T^**W**^out^)^2^] ≥ 0.

The situation is most transparent if we assume that the RNN readout is one-dimensional, in which case the readout and feedback weights become vectors **w**^out^ and b, respectively, and (37) becomes

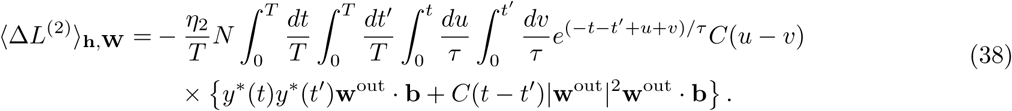

In this case it is clear that, as in the case of feedforward networks [9], the loss function tends to decrease when the readout weights become aligned with the feedback weights. In the following subsection we will show that, at least under similar approximations to the ones made here, such alignment does in fact occur.

### Alignment of readout weights with feedback weights

In the preceding subsection it was shown that, assuming a linear RNN and averaging over activities and recurrent weights, the loss function tends to decrease when the alignment between the readout weights **W**^out^ and the feedback weights **B** becomes positive. Does such alignment indeed occur?

In order to address this question, we consider the quantity Tr(**W**^out^**B**) and ask how it changes following one cycle of training, with combined weight updates on **W** and **W**^out^. (As in the preceding subsection, external input to the RNN is ignored here for simplicity.) The effect of modifying the readout weights is obvious from (13):

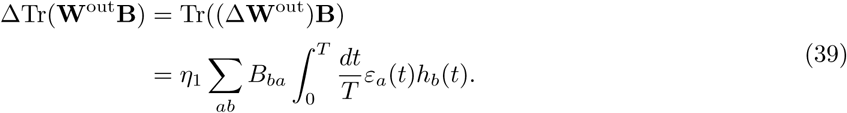

The update to the recurrent weights, on the other hand, modifies **h**(*t*) in the above equation. Because we are interested in the combined effect of the two weight updates and are free to make the learning rates arbitrarily small, we focus on the following quantity:

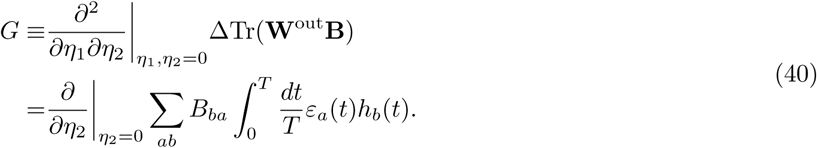

The goal of this subsection is thus to show that (at least on average) *G* > 0.

In order to evaluate this quantity, we need to know how the RNN activity **h**(*t*) depends on the weight modification Δ**W**. As in the preceding subsection, we will assume a linear RNN and will work in the continuous-time limit (*τ* ≫ 1) for convenience. In this case, the dynamics are given by

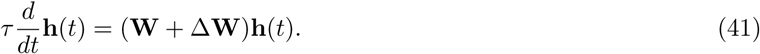

If we wish to integrate this equation to get **h**(*t*) and expand to leading order in Δ**W**, care must be taken due to the fact that W and ΔW are non-commuting matrices. Taking a cue from perturbation theory in quantum mechanics [41], we can work in the “interaction picture” and obtain

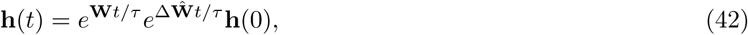

where

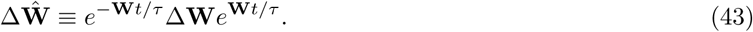

We can now expand (42) to obtain

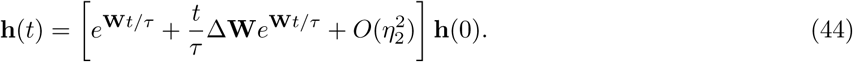

For a linear network, the update rule for **W** from (13) is then simply

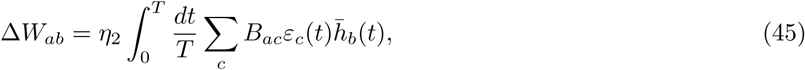

where the bar denotes low-pass filtering:

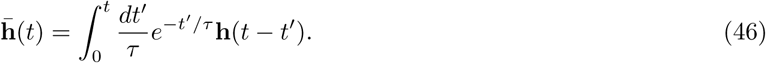

Combining (44)-(46), the time-dependent activity vector to leading order in *η*_2_ is

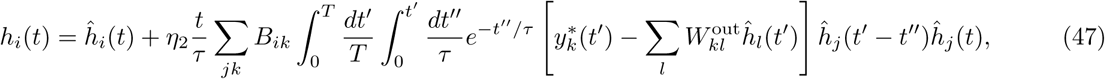

where 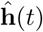 is the unperturbed RNN activity vector (i.e. without the weight update Δ**W**). With this result, we can express (40) as *G* = *G*_1_ + *G*_2_, where

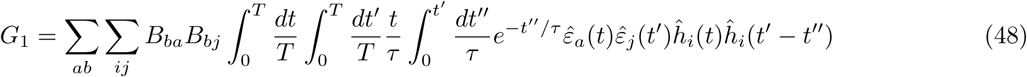

and

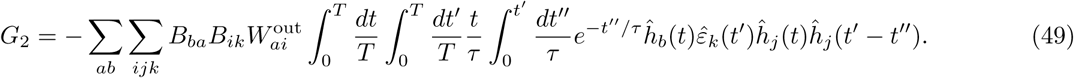

Here we have defined 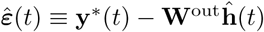.

In order to make further progress, we follow the approach of the previous subsection and perform an average over RNN activity vectors, which yields

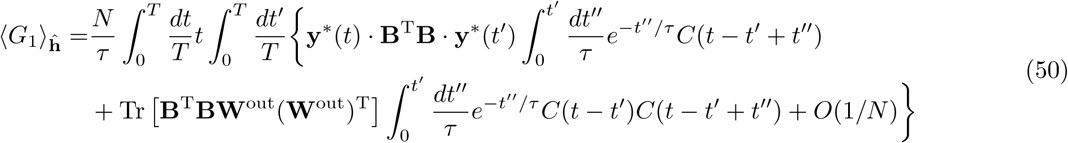

and

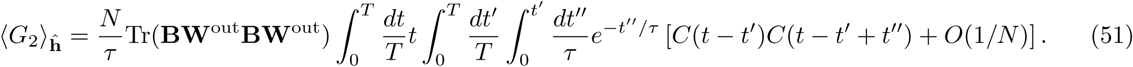

Similar to the integral in (37), both of these quantities will tend to be positive if we assume that *C*(*t*) ≥ 0 with a peak at *t* = 0, and note that the integrand is large only when *t* ≈ *t′*.

In order to make the result even more transparent, we can again consider the case of a one-dimensional readout, in which case (50) becomes

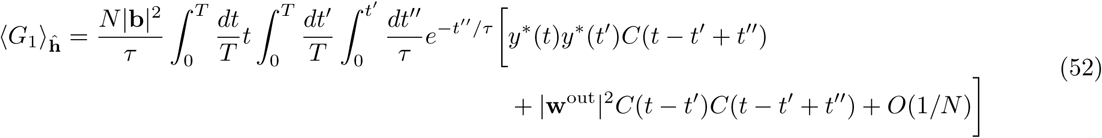

and

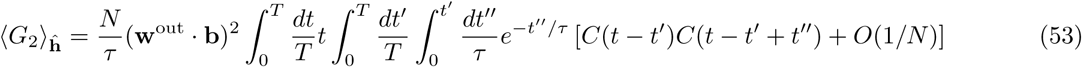

This version illustrates even more clearly that, given a well-behaved target output *y*\*(*t*), the right hand side of this equation tends to be positive.

Equation (50) (or, in the case of one-dimensional readout, (52)) shows that the overlap between the readout weights and feedback weights tends to increase with training. Equation (37) (or (38)) then shows that the readout error will tend to decrease during training given that this overlap is positive. While these mathematical results provide a compelling plausibility argument for the efficacy of RFLO learning, it is important to recall that some limiting assumptions were required in order to obtain them. Specifically, we assumed linearity of the RNN and vanishing of the cross-correlations in the RNN activity, neither of which is strictly true in a trained nonlinear network. In order to show that learning remains effective even without these limitations, we must turn to numerical simulations such as those performed in the main text.

